# Transcription of damage-induced RNA in *Arabidopsis* was frequently initiated from DSB loci within the genic regions

**DOI:** 10.1101/2024.03.15.585142

**Authors:** Kohei Kawaguchi, Soichirou Satoh, Junichi Obokata

## Abstract

DNA double-strand breaks (DSBs) are the most severe DNA lesions and need to be removed immediately to prevent loss of genomic information. Recently, it has been revealed that DSBs induce *de novo* transcription from the cleavage sites in various species, resulting in RNAs being referred to as damage-induced RNAs (diRNAs). While diRNA synthesis is an early event in the DNA damage response and plays an essential role in DSB repair activation, the location where diRNAs are newly generated in plant remains unclear, as does their transcriptional mechanism. Here, we performed RNA sequencing of diRNAs that emerged around all DSB loci in *Arabidopsis thaliana* under the expression of the exogenous restriction enzyme *Sbf* I and observed 88 diRNAs in 360 DSB loci. Most of the detected diRNAs originated within the active genes and were transcribed from DSBs in a bidirectional manner. Furthermore, we found that diRNA elongation tends to terminate at the boundary of an endogenous gene located near DSB loci. Our results provide reliable evidence for understanding the importance of *de novo* transcription at DSBs and show that diRNA is a crucial factor for successful DSB repair.

## 1. Introduction

DNA double-strand breaks (DSBs) are one of the most deleterious types of DNA damage and need to be removed immediately to preserve genomic integrity. Eukaryotes have developed elaborate and complex DSB repair machinery to detect and restore the cleaved DNAs in a precise manner. Recently, it has been revealed that DSBs induce *de novo* transcription near the cleavage sites in various species (Wei et al., 2012; Michelini et al., 2017; Vítor et al., 2019; Böttcher et al., 2022). These RNAs are referred to as damage-induced RNAs (diRNAs), and their synthesis is an early event in the DNA damage response. They contribute to DSB repair activation in various eukaryotes (Wei et al., 2012; Michelini et al., 2017; Bonath et al., 2018; Sharma et al., 2021). In human cells, upon DSB formation, RNA polymerase II (RNAP II) is recruited for diRNA transcription, thereby diRNAs are bidirectionally transcribed from the cleavage sites (Pappas et al., 2023). Transcription of diRNAs is often active in the endogenous genic regions and can be elongated within part or all of the gene body (Aymard et al., 2014; Domingo-Prim et al., 2020). These diRNAs hybridize to damaged double-stranded DNA and form three-stranded nucleic acid structures, designated R-loops, to enhance the efficiency of error-free DNA repair (Yasuhara et al., 2018; Ouyang et al., 2021). Such transcription manner and DSB repair-related function of diRNAs have also been observed to some extent in *Drosophila* and yeast, respectively (Böttcher et al., 2022; Ohle et al., 2016), suggesting that the transcriptional mechanisms and function of diRNAs are highly conserved among eukaryotes. Meanwhile, in plants, the mechanisms by which diRNAs are transcribed, especially in terms of the regulation of their activity and transcribed region, remain unclear.

In this study, we utilized transgenic *A. thaliana* plants harboring the gene for the restriction enzyme *Sbf* I under the control of an inducible promoter (Kawaguchi et al., 2024), and investigated diRNAs at all *Sbf* I-dependent DSB loci using RNA sequencing (RNA-seq) analysis. As a result, we successfully detected *de novo* diRNAs at a variety of DSB loci, and further characterized their transcriptional activity and transcribed regions. Our results provide reliable evidence for understanding the importance of *de novo* transcription at DSB loci and strongly suggest that diRNA is a crucial factor for promoting activation of the DSB repair machinery.

## 2. Results

### 2.1 Genomic distribution and transcriptional competence of diRNA loci

To determine the transcriptional activity of diRNAs on each DSB locus, we used *Sbf* I heat-inducible plant that enables well-controlled DSB induction at multiple *Sbf* I recognition sites (*Sbf* I RS) dispersed throughout the *Arabidopsis* nuclear genome (Kawaguchi et al., 2024). The expression of *Sbf* I gene in this plant could be easily and quickly regulated by simple heat shock treatment (Figure S1). We previously revealed that 360 out of 623 *Sbf* I RS in the *Arabidopsis* genome significantly showed DSB formation upon *Sbf* I digestion (Kawaguchi et al., 2024). For these DSB loci, we performed transcriptomic analysis by RNA-seq and comprehensively investigated whether diRNAs occurred after DSB induction. In this analysis, we defined the transcripts observed only around DSB loci in the *Sbf* I-inducible plants after DSB induction as *de novo* transcripts. We successfully detected novel transcripts at 131 sites; 88 were determined as diRNAs caused by *Sbf* I digestion, and 43 depended on heat-responsiveness (Figure 1a, Table S1). In addition, we classified the genetic context of each cleavage site into four groups [Gene, Intergenic, Transposable element (TE), and Others] based on the TAIR10 genomic annotations (https://www.arabidopsis.org). Interestingly, almost all diRNAs were newly transcribed from *Sbf* I RS within the genic regions, while they were hardly detected in the transcriptionally repressed regions such as the intergenic regions and TEs (Figure 1a, S2). We further investigated the relationship between the transcriptional state of each endogenous gene and diRNA production activity, and found that gene expression levels in diRNA regions were significantly higher than those in non-diRNA regions (Figure 1b). Next, to further elucidate the features of diRNA transcription, we presented averaged profiles around 88 diRNA and 229 non-diRNA loci. We found that the transcription of diRNAs expanded to approximately 1 kb around DSBs, and most of them were deactivated to their baseline levels after 48 hours (Figure 1c, d). These results indicated that DSBs, especially within the active genes, often induced diRNAs in the plant genome.

**FIGURE 1.**
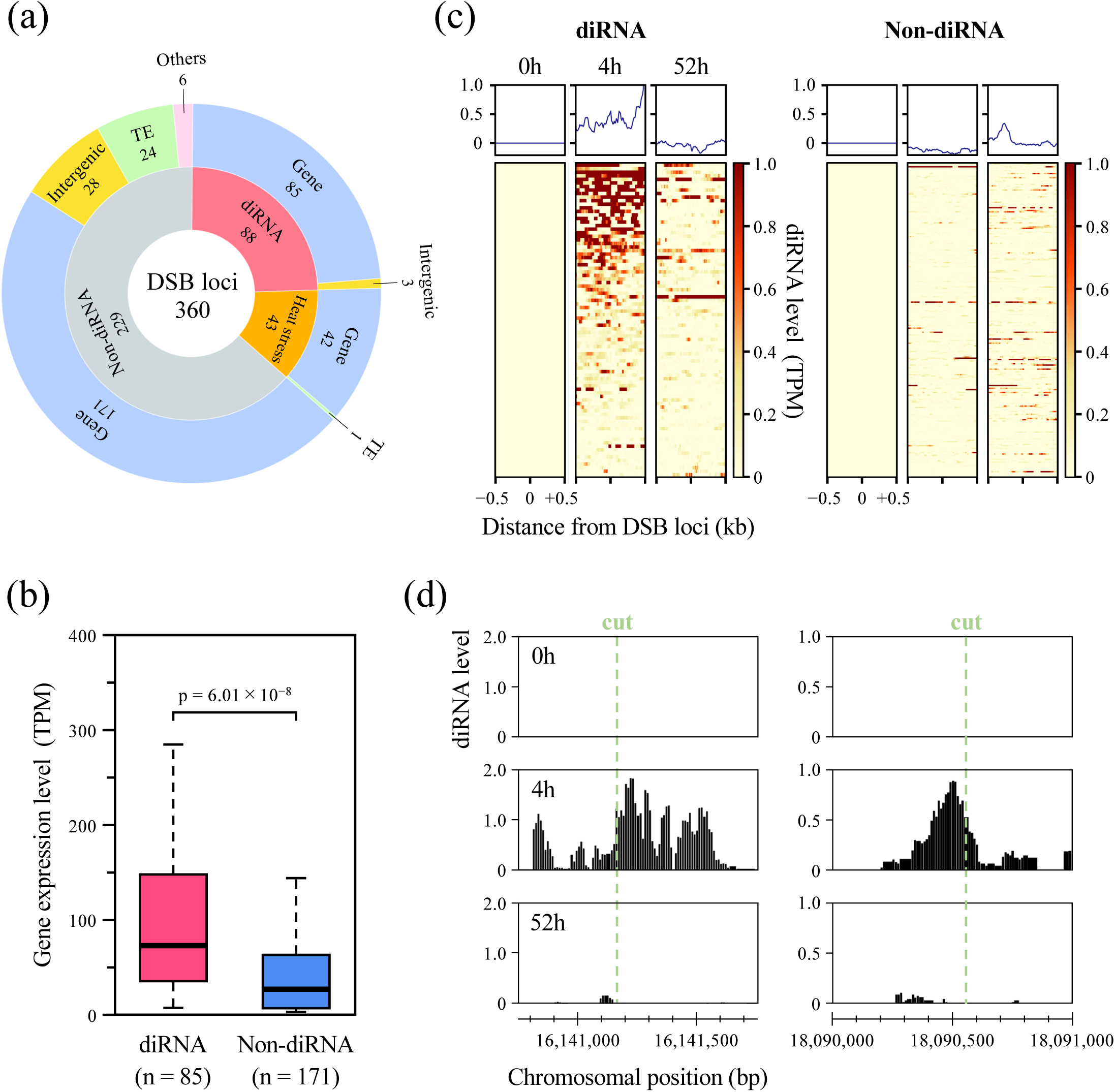
Genome-wide identification of diRNAs at *Sbf* I RS after DSB induction. (a) A sunburst plot showing the number of diRNAs at 360 DSB loci. The inner annulus represents the number of novel transcripts derived from DSBs excluding genes or transposable elements (TEs) caused by heat stress response, and the outer annulus shows the results of classification of each cleavage site based on annotation into four categories (Gene, Intergenic, TE and Others). (b) Boxplots representing gene expression levels of 85 diRNAs and 171 non-diRNAs containing *Sbf* I RS. Wilcoxon rank sum test was used to calculate p-values (p < 0.005). (c) Heatmaps and averaged profiles of *de novo* transcriptions between 88 diRNA and 229 non-diRNA regions. (d) Genome browser screenshots representing transcription levels of diRNAs around DSB loci on chromosome 4.

### 2.2 The majority of diRNAs were transcribed bidirectionally from both cleaved DNA ends

Recently, it has been reported that diRNAs appear to be transcribed in a bidirectional manner from DSB ends in mammals (Michelini et al., 2017; Vitor et al., 2019; Rzeszutek & Betlej 2020). However, it is largely unknown whether this bidirectional manner is the same in other organisms and how often diRNAs initiate transcription induction bidirectionally because insufficient observations have been performed. Therefore, we classified 88 diRNAs into two transcription types: “Bidirectional” and “Unidirectional” (Figure 2a, Table S2). Consistent with previous findings, the majority of diRNAs were newly transcribed from both DSB ends and these levels were approximately equal. Meanwhile, a few diRNAs actively expanded in a unidirectional manner (Figure 2b, c). From these results, we demonstrated that diRNA transcription largely exhibits a bidirectional pattern in *Arabidopsis* and suggested that such transcriptional behavior of diRNAs is widely conserved in various eukaryotes.

**FIGURE 2.**
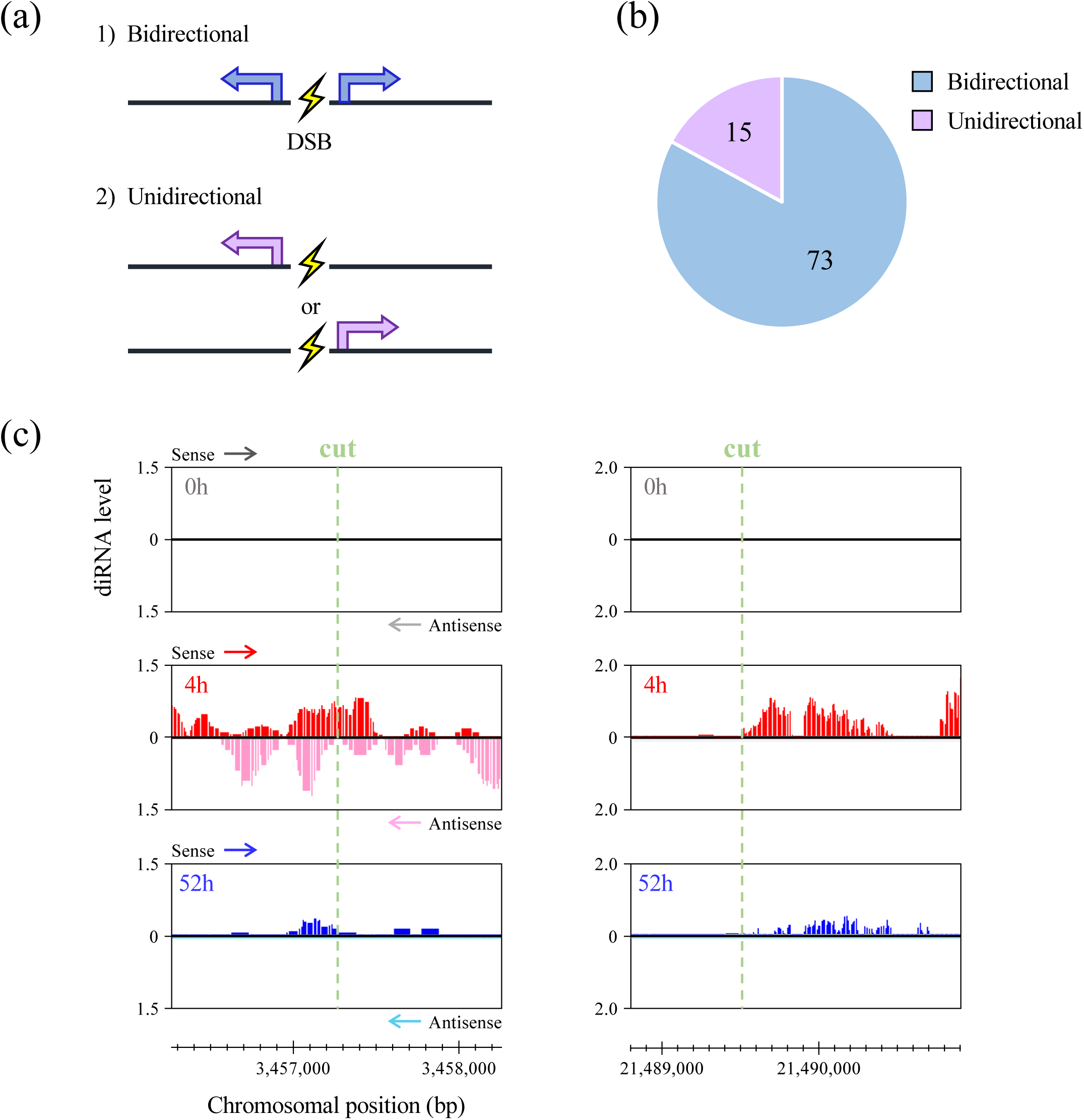
Directional selectivity of diRNA transcription in plants. (a) Two hypothetical models of diRNA transcription at DSBs. (b) Number of diRNAs categorized according to Figure 2a. (c) Genome browser screenshots of bidirectional (left panel) and unidirectional (right panel) diRNA transcription on chromosome 2 and 3, respectively. The orientation of the sense reads is consistent with that of endogenous genes containing the cleavage sites.

### 2.3 Transcription of diRNAs appears to be terminated by sequence motifs similar to those involved in transcriptional termination of protein-coding genes

Finally, we investigated how diRNA transcription is terminated. Surprisingly, some diRNAs detected by RNA-seq in this study exhibited a drastic depletion of mapped reads in close proximity to the transcription start site (TSS) and transcription end site (TES) of pre-existing genes (Figure 3a). We also found that transcription of diRNAs in 35 of the 85 diRNA sites located in genic regions was terminated at the TSS or TES, acting as a border zone (Figure 3b). Moreover, because only poly(A)-tethered RNAs were used for the RNA-seq analysis in this study, the detected diRNAs were assumed to be transcribed by RNAP II (see experimental procedures). From the above, we hypothesized that diRNA transcription might be terminated according to specific transcription termination regulatory elements located near the TSS/TES in pre-existing genes. To examine this hypothesis, we performed motif analysis to determine whether 35 selected border-termination genes have a representative polyadenylation signal (PAS) sequence, composed of “AAUAAA” (Wickens & Stephenson 1984; Yang & Doublié 2011), in the vicinity of the TSS/TES. Overall, 22 of the 35 genes on diRNA-transcribed regions contained the complete PAS sequences around the TSS and TES (Figure 3c). We also investigated a single-base mutation variant, “AUUAAA”, which has also been reported to function as a PAS sequences (Wilusz et al., 1989; Beaudoing et al., 2000). 15 genes with these elements were found near the TSS and TES (Figure S3a). Furthermore, we evaluated the features of the sequence context at the TSS/TES of each gene. These regions are AT-rich and the proportions of each base are relatively similar (Figure S3b). However, known poly(A) cleavage sites in the form of the dinucleotides CA or UA, which are conserved among various species of plants and animals (Rothnie 1996; Li & Hunt 1997; Loke et al., 2005), were enriched only around both the TSS and TES (Figure 3d, e). In summary, our results strongly suggest that diRNA transcription termination is conducted by a 3′ end processing step via RNAP II transcription.

**FIGURE 3.**
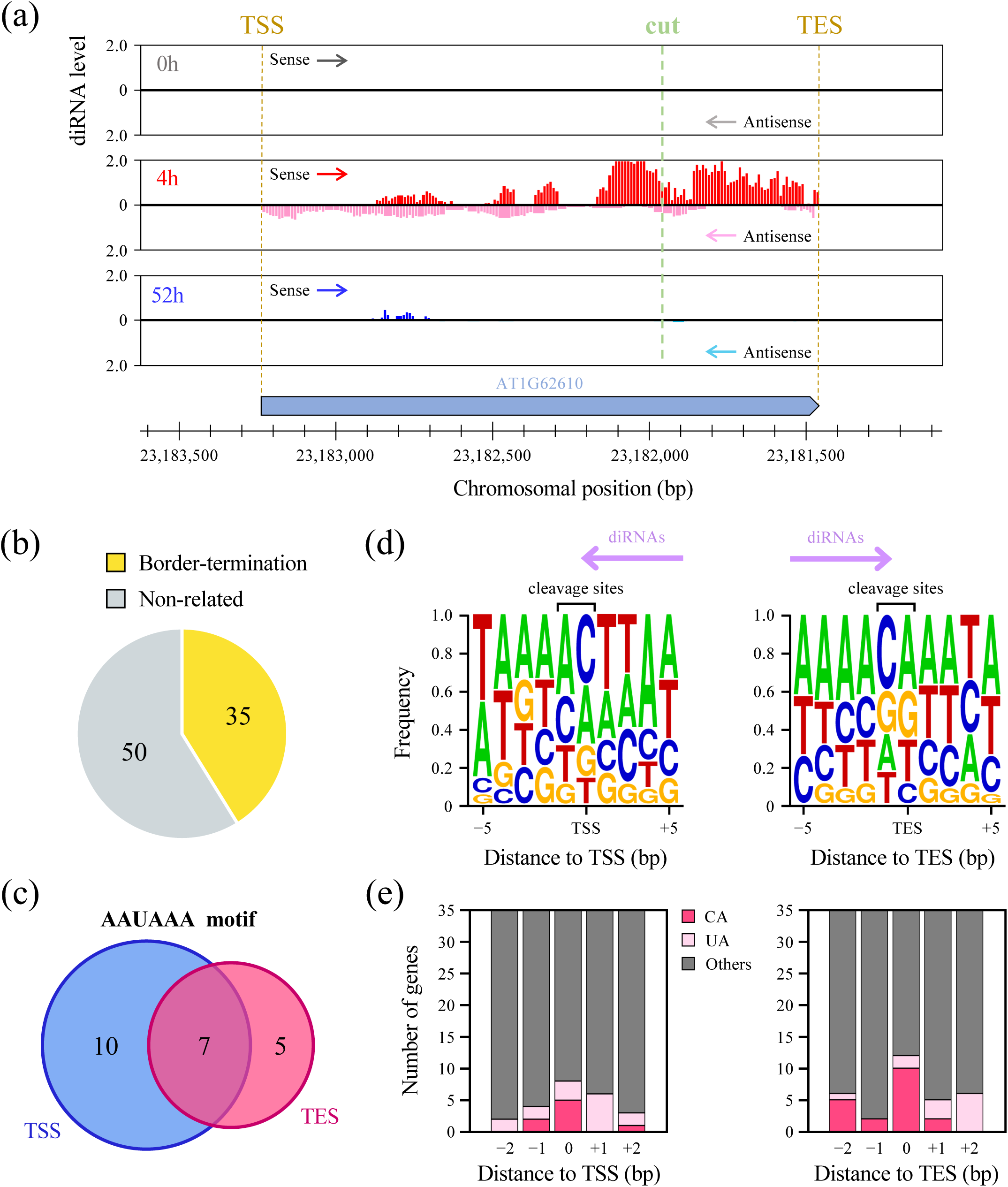
Characterization of diRNA transcription termination occurring within endogenous genes. (a) A genome browser screenshot of diRNA transcription termination near the TSS/TES of AT1G62610. The orientation of the sense reads is consistent with that of endogenous genes containing the cleavage sites. (b) Number of border-termination transcriptomes in 85 diRNAs emerging at genic regions. (c) A Venn diagram representing the number of border-termination genes with poly(A) signals (AAUAAA) within 100 bp of the TSS/TES. (d) The base composition near the TSS/TES in 35 border-termination genes. (e) Number of genes with poly(A) cleavage sites (CA or UA) at the TSS/TES in 35 border-termination genes.

## 3. Discussion

DSB repair is an essential mechanism by which all organisms protect genomic information stored in their DNA from damage. Without effective DSB repair, these breaks accumulate in cells gradually, leading to base mutations and chromosomal rearrangements. Recently, it has been revealed that newly transcribed RNA induced by DSB can interact with exposed DNA ends and facilitates the repair process; this brand-new role of RNA molecules opens up another dimension in the field of DNA repair (Bader et al., 2020; Domingo-Prim et al., 2020). In this study, we detected 88 diRNA loci in the *Arabidopsis* genome (Figure 1a) and found their properties useful for elucidating the activity and initiation/termination mechanisms of diRNA transcription.

First, diRNAs were remarkably transcribed from genic regions with high transcriptional activity (Figure 1b), and rarely observed from the intergenic regions and TEs (Figure 1a). At transcriptionally active genomic regions, RNAP II transcribes vigorously, and its local abundance and turnover rate are very high (Nikolov & Burley 1997; Shilatifard et al., 2003). In contrast, the transcription of TEs and intergenic regions is suppressed by epigenetic regulatory mechanisms such as DNA methylation, histone modifications, and RNA interference (Cui & Cao 2014; Pikaard & Mittelsten Scheid 2014). These epigenetic mechanisms prevent the recruitment of RNAP II onto those genomic regions. Thus, such regulatory mechanisms for RNAP II-related transcription might also be involved in the higher activity of diRNA production at the actively transcribed genic regions. Additionally, diRNAs derived from DSBs play important roles in stimulating the recruitment of homologous recombination (HR) factors for precise DSB repair in eukaryotes (Gao et al., 2014; Rzeszutek & Betlej 2020). Therefore, it is considered that diRNAs in the plant genome also protect the genetic information of active genes from various types of DNA damage.

Next, we uncovered that diRNA transcription initiation and termination share certain features in *Arabidopsis thaliana*. In mammals, diRNAs are generally transcribed in both directions from cleaved DNA ends (Michelini et al., 2017; Vitor et al., 2019; Rzeszutek & Betlej 2020). Additionally, in yeast cells, diRNA transcription elongates within either part or all of a gene when transcription is initiated in the genic region (Aymard et al., 2014; Domingo-Prim et al., 2020). Similarly, we demonstrated comparable properties of diRNA transcription in the plant genome, where bidirectional transcription was observed in almost all diRNA loci (Figure 2), and transcription termination depended on PAS motifs and poly(A) cleavage sites near the TSS/TES of the endogenous genes (Figure 3). It is reasonable that such bidirectional transcription initiation of diRNAs occurs as it would facilitate interaction with both DNA ends for proper DSB repair. Recently, it has been reported that the clearance of newly transcribed diRNA is required for well-controlled DNA end resection and assembly of the HR repair machinery (Domingo-Prim et al., 2019; Zhao et al., 2020; Ngo et al., 2021). On the basis of our results and these findings, the mechanism of initiation of eukaryotic diRNA transcription has important implications for the fidelity of the DSB repair process and this phenomenon is highly conserved across distantly related species. In addition, our analysis exhibited significant signals at multiple cleavage sites, concluding that we observed bidirectional transcription of diRNAs occurring from both damaged DNA ends. However, we cannot completely exclude the possibility that a part of these signals are unidirectional transcripts crossing over DSB loci. In previous study, we suggested that DSBs induced localizations of transcription-related histone marks (Kawaguchi et al., 2024). Such DSB-dependent chromatin remodeling could result in transcriptionally active states of the repaired genes. It has also been indicated that diRNAs can be transcribed in both sense and antisense orientations in mammalian cells (Sharma et al., 2021), consistent with our results. However, as we could not separate the diRNAs which transcribed from DSB sites and passing over repaired DSB sites from around the TSS. To address this issue, full-length sequencing of isoforms of diRNA is necessarily. Meanwhile, regarding the termination of RNAP II-dependent transcription, there are common and specific types of sequence motifs are utilized in plants and animals. Notably, the “AAUAAA” element, which is highly conserved in mammals, acts as a minor signal in *Arabidopsis* genes (Loke et al., 2005). Conventional studies have revealed that poly(A) signals in plants are composed of far upstream elements (FUEs), near upstream elements (NUEs), and poly(A) cleavage sites (Hunt 1994; Li & Hunt 1995; Rothnie 1996; Li & Hunt 1997; Loke et al., 2005). In this study, we found that the 3′ end position of *Arabidopsis* diRNAs was located near both plant and animal types of termination signals. Therefore, it is considered that the fundamental mechanism behind not only the initiation of diRNA transcription but also its termination is common among eukaryotes.

In conclusion, we identified various diRNA transcription events in the plant genome and elucidated their behavior, which involves diRNA production spreading out throughout the genic region from DSB sites. This transcriptional pattern suggests possibilities for novel interactions between RNAP II and DSB repair factors, as well as the contribution of diRNA-dependent DSB repair to the conservation of gene sequences during eukaryotic genome evolution. Extensive future analyses across different plant species and other organisms should deepen our understanding of the mechanisms and significance of diRNA transcription.

## 4. Experimental procedures

### 4.1 Plant materials and growth conditions

*Arabidopsis thaliana* seeds were stratified at 4°C in the dark for 2 days, and then grown on MS medium (Murashige & Skoog, 1962) under continuous light (30–50 μmol m^−2^ s^−1^) at 23°C. 10-day-old seedlings were harvested and subjected to RNA isolation.

### 4.2 Heat-inducible expression of *Sbf* I gene in *Arabidopsis thaliana*

*Sbf* I-inducible plants used in a previous study (Kawaguchi et al., 2024) underwent heat shock treatment at 37°C for 4 hours in an NK-type artificial weather chamber (Nippon Medical and Chemical Instruments Co., Ltd.). After heat induction, all treated plants were subjected to RNA sequencing.

### 4.3 RNA isolation and sequencing (RNA-seq)

Total RNA was isolated from approximately 10–20 seedlings using the RNeasy Plant Mini Kit (QIAGEN) followed by DNase I treatment. After a quality control (QC) check by 2100 Bioanalyzer (Agilent Technologies), RNA-seq libraries were constructed from qualified RNA samples using TruSeq Stranded mRNA Library Prep Kit (Illumina). RNA-seq was performed on an Illumina Nova Seq 6000 platform with a 101 bp paired-end protocol.

### 4.4 RNA-seq data processing

The quality of each set of RNA-seq data is summarized in Table S5. Adapter sequences and low-quality reads were removed by Trimmomatic (ver. 0.39; Bolger et al., 2014) with the following parameters: ILLUMINACLIP:2:30:10 LEADING:20 TRAILING:20 SLIDINGWINDOW:4:15 MINLEN:50. All files were mapped to the reference *A*. *thaliana* genome (TAIR10; https://www.arabidopsis.org/) using STAR (version: 2.7.10a; Dobin et al., 2013) with the following parameters: STAR --alignIntronMin 10 --alignIntronMax 5000, and were processed using SAMtools (ver. 1.6; Li et al., 2009) for the removal of PCR-based duplicated reads (rmdup). Coverage calculations and transcripts per kilobase million (TPM) normalization of each aligned RNA-seq dataset (BAM files) were performed in deepTools (ver. 3.5.2; Ramírez et al., 2014) with the following parameters: bamCompare -of bigwig --binSize 10 --scaleFactorsMethod None --normalizeUsing BPM.

### 4.5 Genome-wide diRNA detection using RNA-seq data

First, we calculated the total coverage of 500 bp around 360 DSB loci using a Perl script, followed by subtracting (1) the value at 0h from that at 4h, and (2) the value at 52h from that at 4h for each *Sbf* I RS. If the two values after the calculation of (1) and (2) exceeded 0, such *Sbf* I RS was regarded as a novel transcription after heat shock treatment. Moreover, among these *Sbf* I RS, DSB loci located at heat stress-responsive genes/TEs as indicated in *Arabidopsis* eFP browser (Winter et al., 2007) were excluded. In addition, a heatmap and averaged profile surrounding DSB loci were generated from the obtained matrix data files using deepTools (computeMatrix and plotHeatmap tools).

### 4.6 Motif analysis

Using FIMO (Grant et al., 2011), we searched the specific poly(A) motifs (AAUAAA and AUUAAA) present in the region 50 bp upstream and downstream of the TSS and TES. Motif analysis data are listed in Table S4. Sequence logo plots around the TSS and TES were generated by WebLogo (Crooks et al., 2004).

## Author contributions

K. Kawaguchi, S. Soichirou, and J. Obokata designed this research; K. Kawaguchi performed the biological experiments and analyzed the data; K. Kawaguchi and S. Soichirou wrote the manuscript; and K. Kawaguchi, S. Soichirou, and J. Obokata performed a proofreading of the manuscript. All authors read and approved the final manuscript.

## Acknowledgements

We thank M. Kazama, K. Mukae, Y. Shigematsu, S. Morita and H. Narukawa for experimental supports, discussions and encouragement. This work was supported by Japan Society for the Promotion of Science (JSPS) KAKENHI (22K06342 to S.S.) and Kyoto Prefectural University Academic Promotion Fund Research Encouragement Program (SK20221024 to K.K.).

## Data Availability

The data underlying this article are available in the article and in its online supplementary material. RNA-seq data have been deposited to the DNA Data Bank of Japan (DDBJ) database under the accession codes DRA018155 (DRR537390−DRR537392).

## Supplementary figure legends

**Figure S1.**
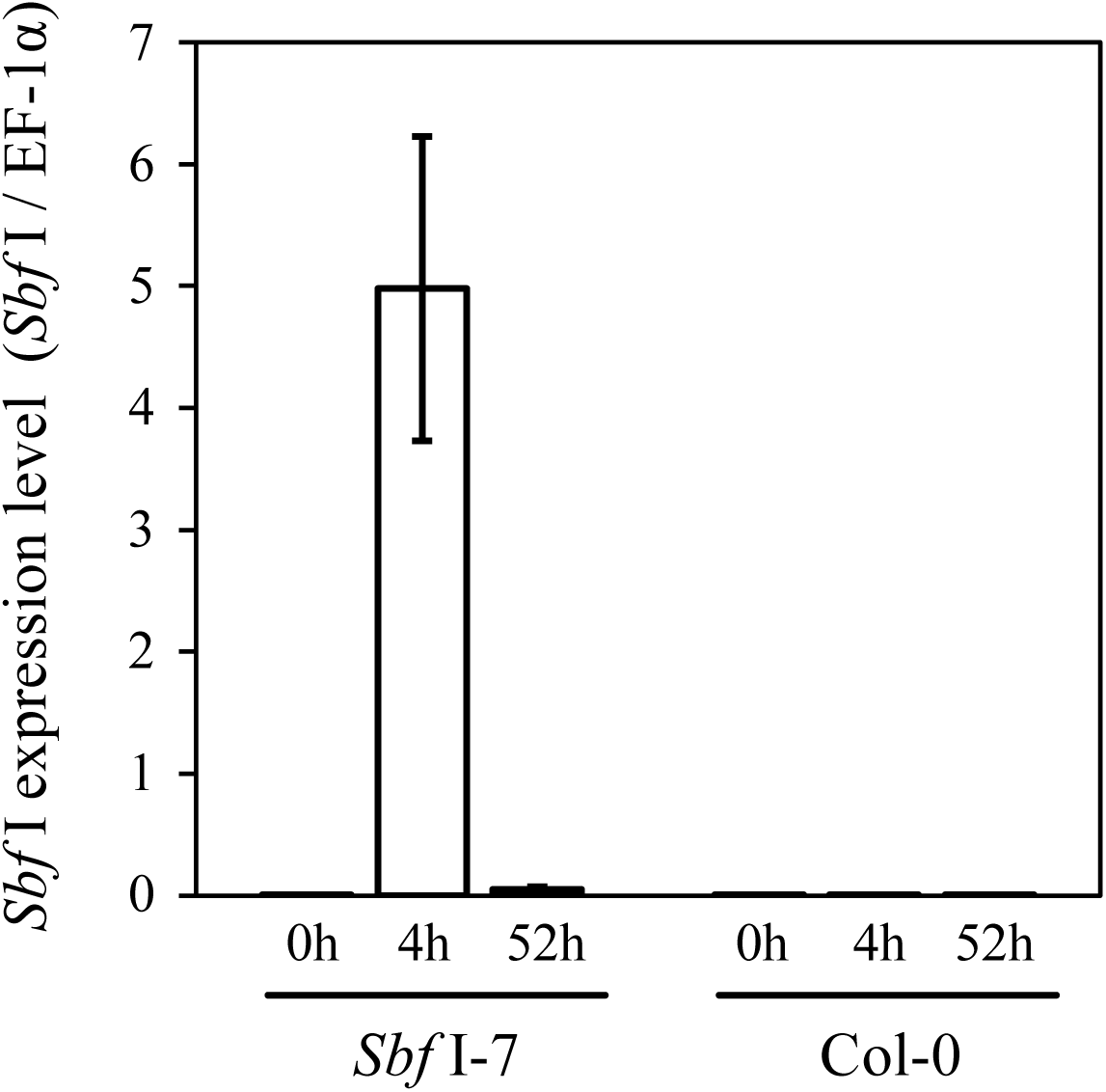
Validation of *Sbf* I transcription level after HS treatment by RT-qPCR. Extracted total RNA was reverse-transcribed using random hexamer, followed by use of the resultant cDNA as templates for expression analysis by RT-qPCR. The PCR primer sets are described in Table S6. Relative RNA levels of the *Sbf* I gene were calculated and normalized to internal controls, the AT5G60390 (EF-1α) gene-encoded GTP-binding elongation factor (Czechowski et al., 2005; Udvardi et al., 2008). Col-0 (wild-type) sample was used as a negative control. Error bars represent ±SD of three biological replicates.

**Figure S2.**
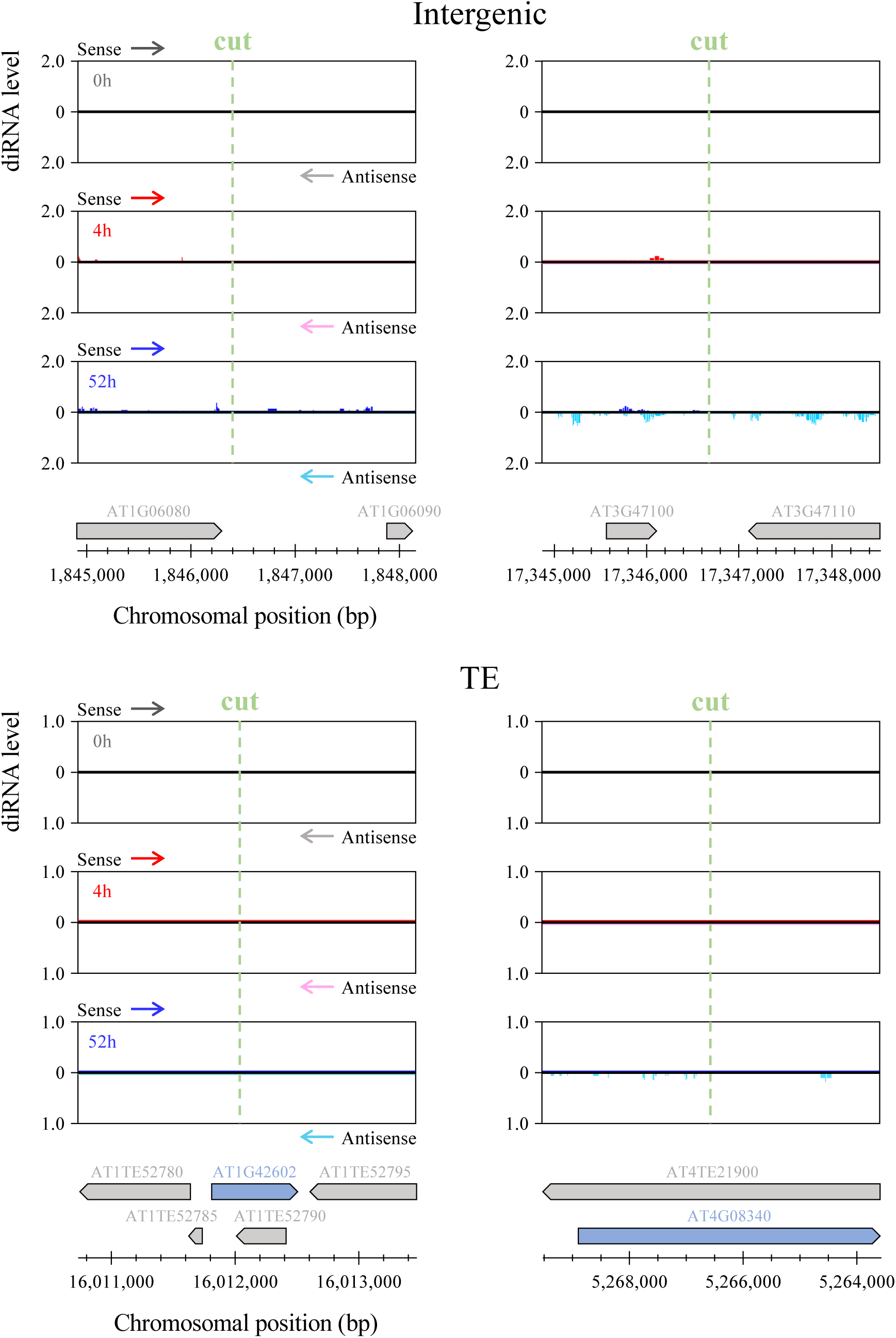
Transcriptional states of non-diRNA regions after DSB induction. Genome browser screenshots representing intergenic (upper panels) and TE (lower panels) regions containing *Sbf* I RS. The orientation of the sense reads is consistent with that of blue endogenous genes containing the cutting sites.

**Figure S3.**
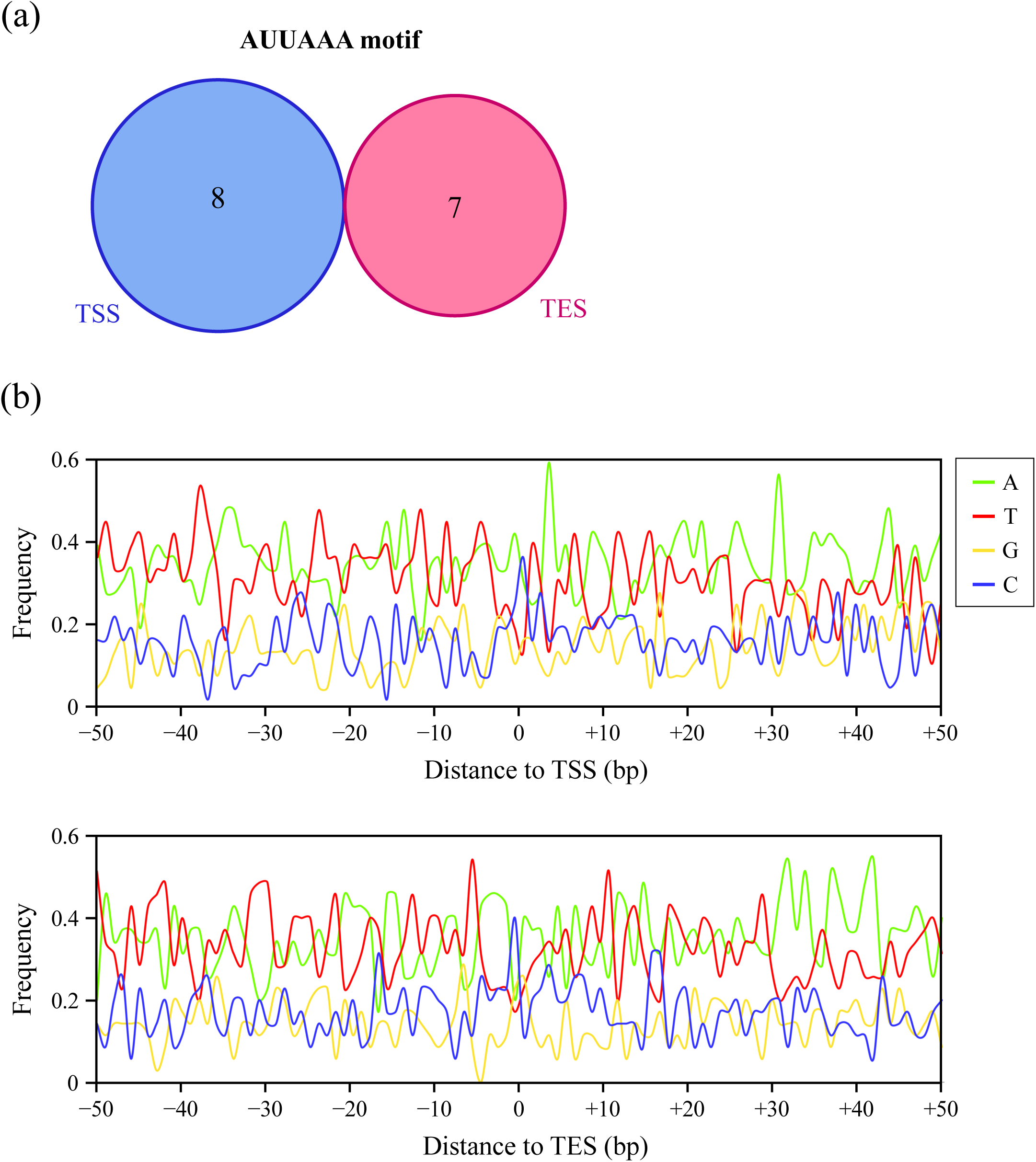
Motif analysis and base compositions of border-termination genes. (a) A Venn diagram representing the number of border-termination genes with poly(A)-like signals (AUUAAA) within 100 bp of TSS/TES. (b) Distribution of A, T, G, and C contents (%). The x- and y-axes show the base position from TSS/TES and the percentage of each base sequence, respectively.

## Supplementary Tables

Table S1: Determination of diRNAs at 360 DSB loci by RNA-seq analysis.

Table S2: Classification of diRNA direction into two types; ’Bidirectional’ or ’Unidirectional’ at 88 *Sbf* I RS.

Table S3: Information of 35 border-termination diRNAs.

Table S4: Motif analysis of AAUAAA and AUUAAA signals within upstream and downstream 50 bp of the TSS and TES for 35 border-teamination diRNAs.

Table S5: Total read counts and mapping statistics of RNA-seq data in *Sbf* I-inducible plants.

Table S6: PCR primers used in study.

